# Computational Pipeline to Identify Gene signatures that Define Cancer Subtypes

**DOI:** 10.1101/2022.11.20.517258

**Authors:** Ekansh Mittal, Vatsal Parikh, Raphael Kirchgaessner

## Abstract

**Motivation:** The heterogeneous nature of cancers with multiple subtypes makes them challenging to treat. However, multi-omics data can be used to identify new therapeutic targets and we established a computational strategy to improve data mining.

**Results:** Using our approach we identified genes and pathways specific to cancer subtypes that can serve as biomarkers and therapeutic targets. Using a TCGA breast cancer dataset we applied the ExtraTreesClassifier dimensionality reduction along with logistic regression to select a subset of genes for model training. Applying hyperparameter tuning, increased the model accuracy up to 92%. Finally, we identified 20 significant genes using differential expression. These targetable genes are associated with various cellular processes that impact cancer progression. We then applied our approach to a glioma dataset and again identified subtype specific targetable genes.

**Conclusion:** Our research indicates a broader applicability of our strategy to identify specific cancer subtypes and targetable pathways for various cancers.

## 1. Introduction

Cancer is one of leading causes of death. Approximately 39.5% of women and men will be diagnosed with cancer at some point during their lifetimes. Over 10 million people are estimated to die from the various forms of cancer in the world each year, as stated by source. Within the United States, in 2022 alone, there will be an estimate of 1.9 million new cancer cases diagnosed and 609,360 deaths will occur due to cancer (Cancer Facts & Figures 2022). Current treatment of cancer involves nonspecific methods such as chemotherapy and radiotherapy, which cause an equivalent amount of damage to the patient’s healthy cells as they do to the cancer cells. However, there is much evidence suggests that precision therapy or personalized medicine for cancer patients could potentially reduce the side effects and toxicity caused of chemotherapy and radiotherapy (Kenneth and Cho, 2022; Manrriquez, et al., 2022; Ronquillo and Lester, 2022). Therefore, recent efforts have been made to implement precision therapy for cancer treatment(Krzyszczyk, et al., 2018; Mateo, et al., 2022). For this, new biomarkers and drug targets need to be identified. The influx of ground-breaking whole-genome DNA sequencing and gene expression technologies has allowed for an increase in the characterization of the cancer genome and transcriptome, allowing for the discovery of new genetic biomarkers (Cancer Genome Atlas Research, et al., 2013; Zhang and Wang, 2015). However, the rapid processing of this vast amount of data for the identification of the most significant and functionally relevant changes has been a challenge for cancer identification and treatments. Therefore, new high-throughput strategies have to be proposed and implemented, such as the use of Machine Learning.

Machine Learning approaches can be used to prioritize the most significant genes associated with specific disease subtypes to determine their clinical significance (Alabi, et al., 2021; Arjmand, et al., 2022; Cai, et al., 2022; Chung, et al., 2022; Guo, et al., 2022; Tian, et al., 2021; Tsai, et al., 2022). Recently, strides have been made to incorporate machine learning approaches into cancer therapy (Malebary and Khan, 2021). However, there are limited approaches which are focused on the identification of subtype specific gene signatures tailored for therapeutic implications. Our goal is to develop a machine learning-based pipeline that can identify novel cancer subtype-specific therapeutic targets based on gene expression data.

In this study, we focused on developing and applying the machine learning approaches for the identification of breast cancer subtypes. Additionally, we developed a workflow to identify the associated signaling mechanisms and therapeutic targets that are specific to each breast cancer subtype. Breast cancer is the most frequently diagnosed cancer and the second cause of cancer deaths in women. In 2022, 287,850 women were estimated to have been diagnosed with invasive breast cancer in the United States and approximately 43,250 women were estimated to have died of their disease (Cancer Facts & Figures 2022). Men can also have breast cancer, although male breast cancer is rare. Despite of all the efforts these estimated numbers are significantly higher than the cases diagnosed in prior years (Cancer Genome Atlas, 2012; Ciriello, et al., 2015; Jiang, et al., 2016; Kalecky, et al., 2020). Therefore, existing methods for therapeutic target identification demand improvement.

Further, we utilized a gliomas dataset to validate our pipeline. Gliomas are some of the most aggressive and mostly lethal forms of brain cancer. They typically affect people above the age of 50 and the standard therapy includes surgery, chemotherapy, and radiotherapy. However, the average survival rate is 14 to 15 months, following the standard therapy. Therefore, better markers and treatments to improve patient outcomes are needed. Our glioma dataset contained three molecular subtypes. Two of the subtypes were low-grade gliomas: oligodendrogliomas and astrocytomas. The third subtype was glioblastomas, which is the most severe form of brain cancer and has a poor prognosis (Zhang, et al., 2019). Here we developed a ML approach that can be used to identify specific subtypes of cancers using breast cancer and gliomas dataset.

## 2. Materials and methods

### 2.1 Expression dataset

We utilized Pan-TCGA data (The Cancer Genome Atlas). The dataset included the expression values of over 8000 genes, 6900 cell samples, and 20 different cancers. The goal was to create a pipeline that would allow for the identification of novel subtype-specific target genes for a given cancer. For a proof of concept, cancer with the highest number of samples, breast cancer, was utilized to train the model and develop the pipeline. This pipeline is then applied to the glioma dataset.

### 2.2 Normalization and encoding of expression dataset

The data was encoded and normalized, so a logistic model could be trained on it. For our data specifically, the x-matrix, gene expressions, was normalized, and the y-matrix, subtype classifications, was encoded with a nominal encoder. For the x-matrix dataset the scikit-learn normalizer was utilized (Bac, et al., 2021) and it allowed for all the gene expression values to be within the range of 0.0 and 1.0 since that is the format the input features need to be for logistic regression (Matplotlib 3.5.1). Furthermore, the y-matrix had to be encoded from strings to numbers, since the y-matrix contained the subtypes, which were stored as such, [“Luminal A”, “Luminal B”, “Triple negative (Basal)”, “HER2+”] (Niklaus, et al., 2021; Romero-Cordoba, et al., 2021). This was thus encoded using an ordinal encoder allowing the strings to be expressed as numbers demonstrated in **Figure 1A**. Once all the data was normalized and encoded it served as input for the logistic regression model.

**Figure 1.**
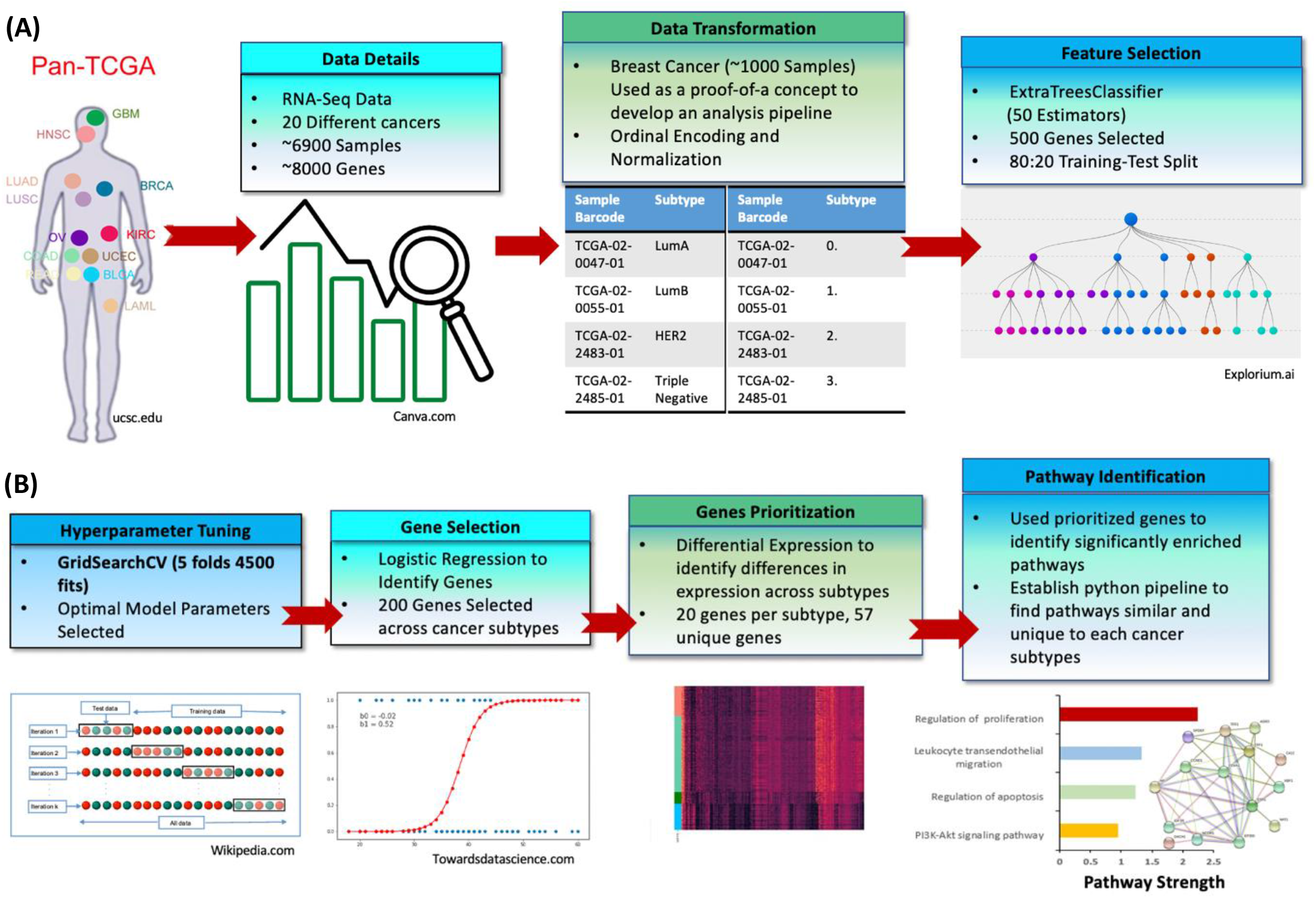
Schematic describing the methods. The work flow describing (A) data extraction, data transformation, and feature selection for machine learning analysis and (B) gene selection, gene prioritization, and pathway analysis

### 2.3 Machine learning analysis

A logistic regression model was utilized for this dataset (Yuan, et al., 2022). There were two different approaches made to train the model. One model was trained without the use of a feature selection method and the other model was trained with the use of a feature selection method, ExtraTreesClassifier (ETC) (Lage, et al., 2022; Lee, et al., 2020). Without the use of the ETC, all 8000 genes were used as input features to train the model; however, once the ETC was implemented the input feature number was brought down to 500 allowing for a much more precise model. The overall model accuracy increased by 7 percent after implementing ETC. Moreover, the precision for predicting each individual subtype after reducing the input features from 8000 to 500 was improved. [RK1] Additionally, to improve model accuracy and ensure the validity of our model, we performed GridsearchCV hyperparameter tuning. GridsearchCV works by splitting the training data into K folds (K=5) and then trains a logistic regression model with hyperparameters defined in an inputted grid on each fold. After training K models for each combination of hyperparameters, the algorithm uses various metrics, such as penalty, solver, C-value, and maximum iterations, to assess the model and hyperparameters, selecting the best ones, therefore optimizing the model. Once the ideal parameters are identified they can be set for the logistic regression model and it can be trained on the data. After the model was trained, the gene’s significance was ranked based on the coefficients they were assigned by the model.

### 2.4 Differential expression

From the 500 input features used to train the model, the top 200 were selected based on the highest coefficients which were assigned by the model based on how predictive the genes were of disease. From those 200 genes, differential expression analysis was used to calculate the p-values and impact factors of all the genes. Having access to these calculations then allowed us to create a comprehensive list of the top 20 most predictive genes for each molecular subtype. All of these genes were then taken and plotted onto a heatmap allowing for relationships to be seen, and the heatmap also allowed for the visualization of the genes that were the distinguishing factors between the different molecular subtypes.

### 2.5 Pathway analysis for gene interaction and associated functions

The protein-protein interaction (PPI) network was constructed using the Search Tool for the Retrieval of Interacting Genes (STRING) database (Szklarczyk, et al., 2015). The STRING tool performs network analysis based on experimental and knowledge-based evidence. The same tool was used to perform functional enrichment analysis of the differentially expressed genes (DEGs), the corresponding biological processes (BP), cell components (CC), and molecular functions (MF) were identified using Gene Ontology (GO), and the signaling pathways involved were identified using the Kyoto Encyclopedia of Genes and Genomes (KEGG) for the *p* adjusted value < 0.05. Pathway strength is defined as the percentage of genes represented in our dataset for a given pathway (**Figure 1B**).

## 3. Results

### 3.1 A hyper-tuned machine learning approach using logistic regression combined with ExtraTreesClassifier (ETC) has better accuracy for cancer subtype prediction than just a logistic regression model

Two different machine learning approaches were applied to train the model. One model was trained using logistic regression without the use of a feature selection and the other model was trained with the use of a feature selection method, ETC, a dimensionality reduction technique. The overall model accuracy increased by 7 percent (79% to 86%) after implementing ETC. The precision for predicting each individual subtype after reducing the input features from 8000 to 500 is improved except for the Triple-Negative subtype (**Figure 2**). However, after conducting hyperparameter tuning the overall predictive accuracy of the model increased to 92%. Additionally, the predictive accuracy for Triple-Negative also increased. Furthermore, throughout the entire model development process, the HER2+ subtype was the most difficult for the model to predict due to the limited number of samples in the data. Overall, the model has very high accuracy, figure 2 also indicates that false positives of Luminal A and B were very rare and only occurrences for HER2+ and Triple-Negative. This makes sense because those two subtypes are molecularly challenging to identify.

**Figure 2.**
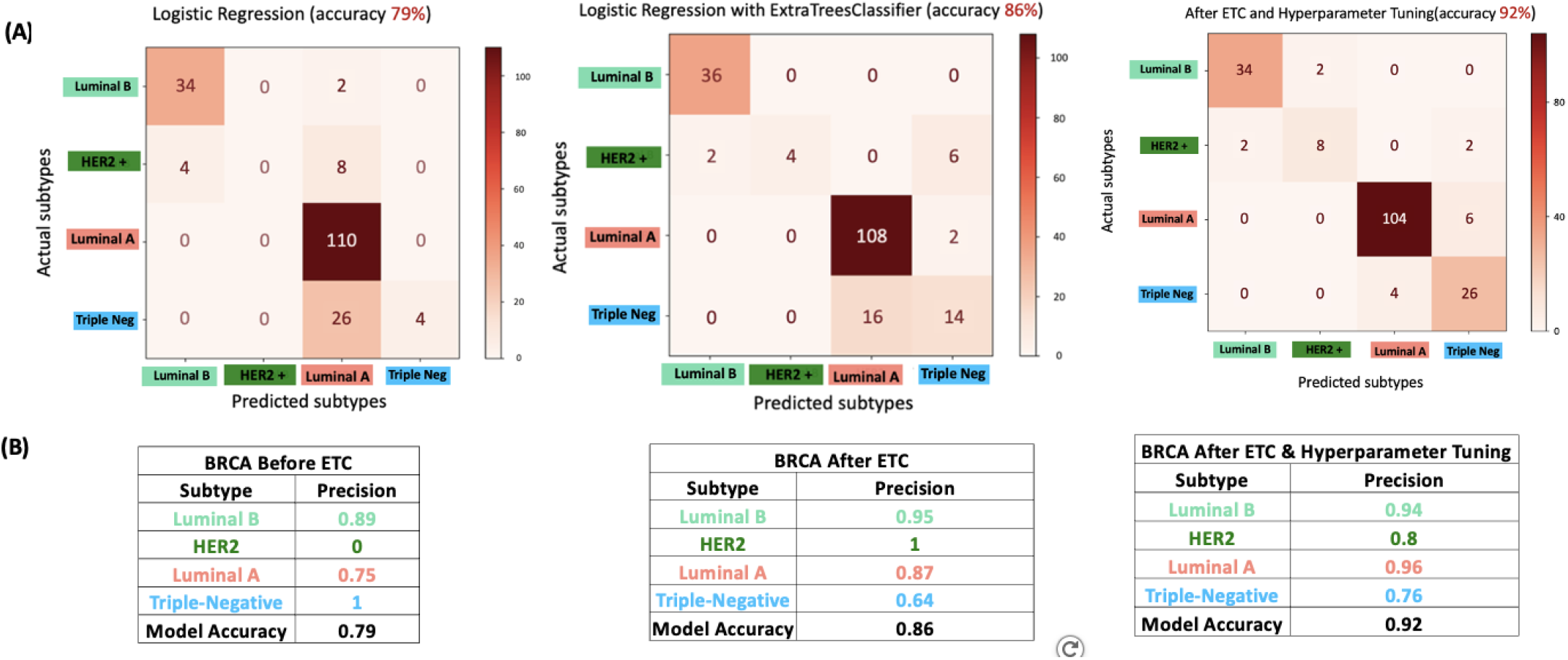
Integrating ExtraTreesClassifier (ETC) and Hyperparameter tuning had improved model accuracy for all the breast cancer subtypes. (A) Confusion Matrix for each breast cancer subtype showing distribution of predictions between subtypes for Logistic Regression alone, Logistic Regression + ETC, and Logistic Regression + ETC + Hyperparameter tuning. (B) Tables showing the precision for each breast cancer subtype for breast cancer with logistic regression alone, Logistic Regression + ETC, and Logistic Regression + ETC + Hyperparameter tuning. The numbers showed include the showing number of trained samples for predicted versus actual subtypes.

### 3.2 Differential expression analysis identified the similarities and differences in each cancer subtype

After the models were optimized, differential expression analysis was used for each breast cancer subtype. From the top 200 genes that were retrieved from the model, 20 genes that had the highest differential expression values were selected. Using these genes, we identified that luminal A and luminal B are closely related while HER2+ and Triple-Negative breast cancer subtypes are most distinct (**Figure 3A**). Furthermore, differential expression allowed for the driving genes to be identified, which could then be further analyzed using protein libraries to examine protein pathways and protein-protein interactions.

**Figure 3.**
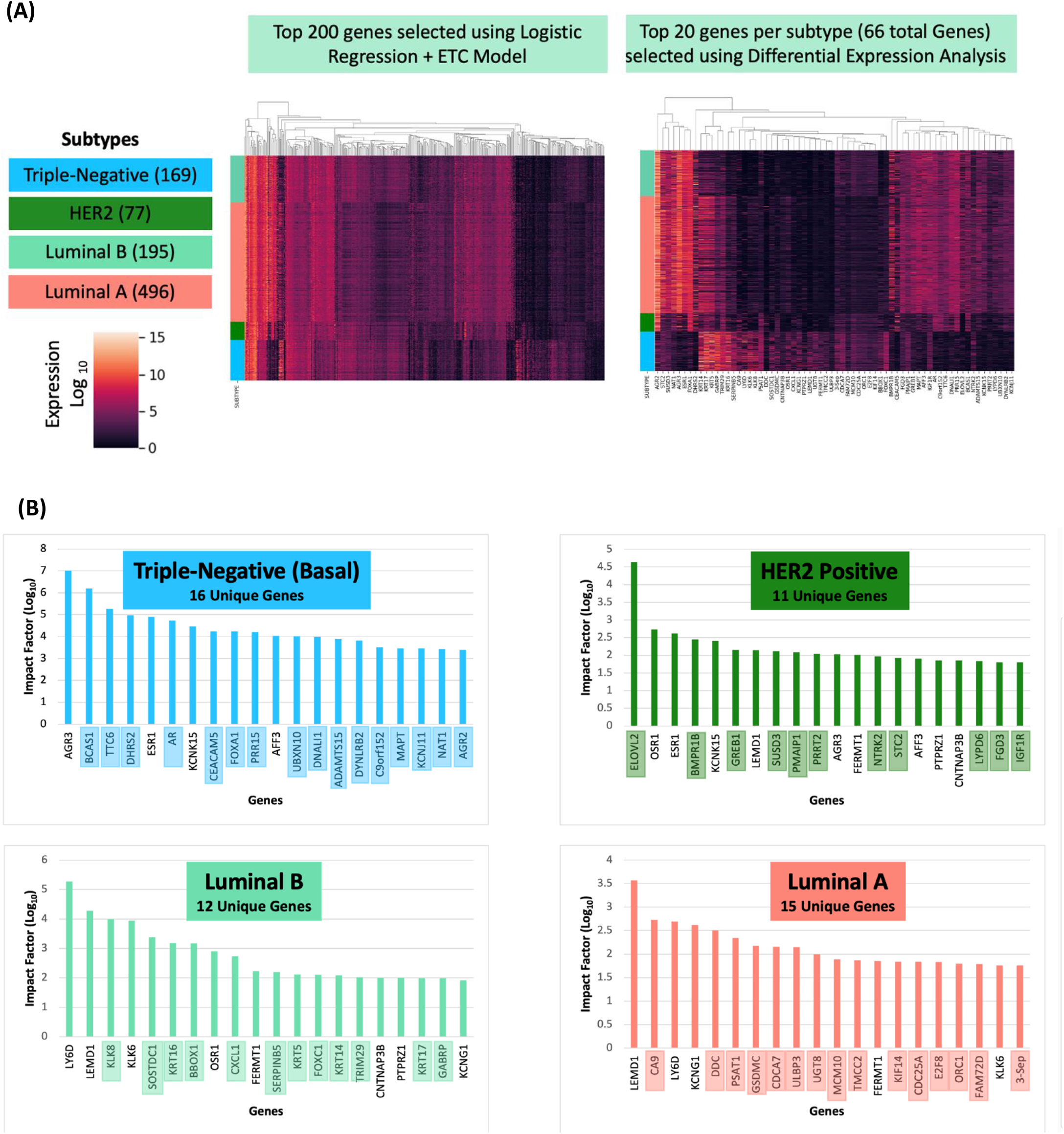
Differentially expressed genes across subtypes of breast cancer. **(A)** The heatmap analysis of differentially expressed genes showing top 200 differentially expressed genes (left panel) and 20 genes per subtype (66 total genes, right panel) selected using logistic regression and ExtraTreesClassifier for each breast cancer subtype. **(B)** Top 20 differentially expressed genes for each breast cancer subtype. Genes which have the highest impact factor and are most significant (less than 0.05 p value) were selected. Impact factor for each gene is defined by the ratio of gene expression (log10) for each cancer subtype over gene expression (log10) for normal breast cancer subtype as defined by Cancer Genome Atlas (TCGA).

### 3.3 Each breast cancer subtype is associated with a distinct gene signature

Analysis of the top 20 differentially expressed genes for each breast cancer subtype identified a distinct gene signature for all 4 subtypes including Luminal A, Luminal B, HER2+, and Triple-negative (BASAL) subtypes. Cross overlap of these genes across these 4 subtypes indicates that there are 15 unique genes associated with Luminal A, 12 unique genes with Luminal B, 11 unique genes with HER2+, and 16 unique genes are associated with Triple-negative breast cancer subtype (**Figure 3B**).

### 3.4 Protein-protein network analysis of genes and therapeutic implications

The network analysis of 20 differentially expressed genes for each breast cancer subtype were analyzed by the STRING pathway analysis for protein-protein interactions. The string analysis curates interactions based on existing experimental and knowledge-based evidence (**Figure 4A**). These 20 genes are associated with various cellular and molecular processes that have high significance. Each breast cancer subtype shows a network cluster specific to the cancer subtype (**Figure 4B**). Luminal A shows a network of LEMD1, CA9, LY6D, KCNG1, DDC, PSAT1, GSDMC, CDCA7, ULBP3, UGT8, MCM10, TMCC2, FERMT1, KIF14, CDC25A, E2F8, ORC1, FAM72D, KLK6, and 3-Sep genes. These genes are associated with the enrichment of cell cycle, proliferation, and DNA damage response pathway. Luminal B shows a network similar to Luminal A but still distinct (LY6D, LEMD1, KLK8, KLK6, SOSTDC1, KRT16, BBOX1, OSR1, CXCL1, FERMT1, SERPINB5, KRT5, FOXC1, KRT14, TRIM29, CNTNAP3B, PTPRZ1, KRT17, GABRP, KCNG1). These genes are enriched in migration, papillomas, keratosis, and cancer development. HER2 network is distinct from both Luminal A and B and only have a few overlapping nodes with Luminal B including OSR1, LEMD1, FERMT1, PTPRZ1, and CNNAP3B. Finally, the network of Triple-negative subtypes is also distinct with only a few similarities with other subtypes including AGR3, ESR1, AFF3, and KCNK15. Additionally, HER2 positive and Triple negative breast cancer subtypes have enrichment of pathways associated with mammary gland development and epithelial cell proliferation (**Figure 4A**). After the identification of network clusters, we performed a literature search to identify if any of these networks are also targetable using small molecules. Interestingly, we identified that for Luminal A, MCM5 (Mini-chromosome maintenance protein complex) can serve as a potential therapeutic target and can be targeted using micro-RNAs (noncoding RNAs that can suppress gene expression) such as miRNA-214 (Wang, et al., 2022). MCM5 is one of the essential components of the pre-replicative complex and forms a helicase together with other proteins to unwind the DNA duplex in S phase. Thus is critical for DNA replication and DNA damage response. For Luminal B, KRT5 (Keratin 5) can be targeted using miR-601(Du, et al., 2019). For HER2 positive, NTRK2 (Neurotrophic Receptor Tyrosine Kinase 2) and IGF1R (insulin-like growth factor type I receptor) are targetable. Both NTRK2 and IGF1R are tyrosine kinase and have role in cell growth and differentiation in cancer and other diseases. NTRK2 can be targeted using entrectinib, which is currently in clinical trial (Wang et al., 2020), and IGF1R can be targated using BMS-754807 and OSI-906 (linsitinib) (Murakami, et al., 2016) (Fuentes-Baile, et al., 2020). Finally, the most aggressive type of breast cancer Triple-negative can be targeted using FOXA1 combination with IGF1R inhibitor (Calissi, et al., 2021) (Calissi et al., 2021) (**Figure 4B**). Overall, our work identified a specific target for each breast cancer subtype. However, more work is needed to validate the function of each target in these cancer subtypes and validate the efficacy of these small molecules.

**Figure 4.**
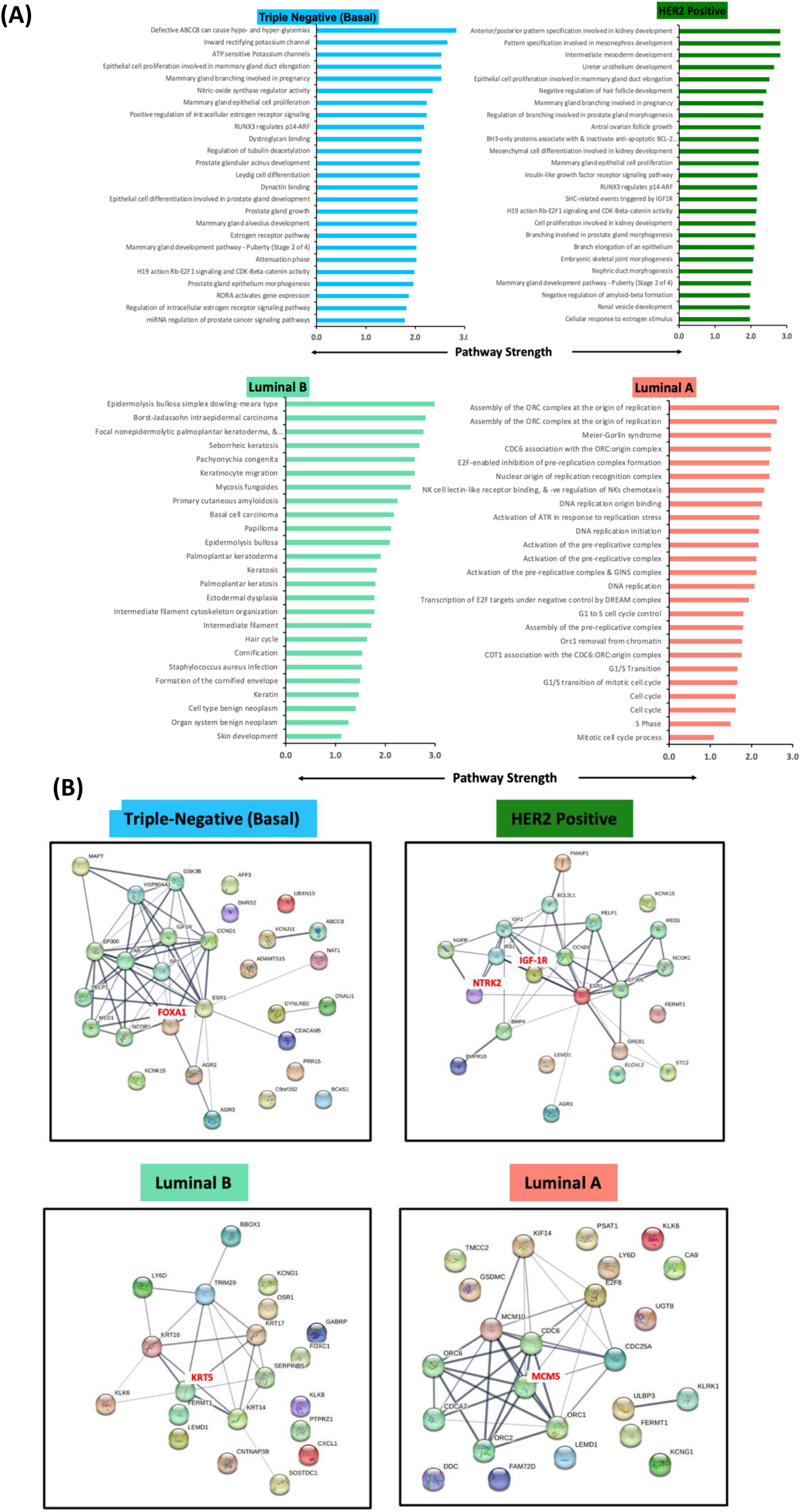
(A) Top 20 differentially expressed genes for each breast cancer subtype was integrated with STRING data analysis. The same tool was used to perform functional enrichment analysis of the differentially expressed genes that are corresponding biological processes (BP) and molecular functions (MF) which were identified using Gene Ontology (GO) and the signaling pathways involved were identified using the Kyoto Encyclopedia of Genes and Genomes (KEGG) for the *p* adjusted value < 0.05 with top 25 for pathway strength. The color coding represents broader categories of these terms. *Pathway strength is defined as percentage of genes represented in our dataset for a given pathway. (B) Protein-protein interaction network analysis of differentially expressed genes for each breast cancer subtype were generated using String-DB and are curated based on string evidence from knowledge and experimental based curations. After the identification of network clusters, we performed a literature search to identify targetable pathways and associated small molecules.

### 3.5 Validation of Computational Pipeline Using Gliomas Dataset

To validate the accuracy of the computational pipeline, a gliomas dataset from TCGA was utilized. Using a logistic regression model with tuned hyperparameters, and ETC, we were able to achieve a subtype predictive accuracy of 96% (Figure 5A). From this model, the top 200 genes for each subtype were further analyzed with differential expression analysis. Through differential expression, we identified the top 20 genes for each subtype (Figure 5B-C), which were then further analyzed using StringDB pathway analysis (Figure 5D). Through the pathway analysis, we identified matrix metalloproteinases (MMP3) can potentially be targeted in glioblastomas (GBM). MMP3 over-expression has been associated with cancer metastasis and tumor growth in various cancers including breast cancer (Liang, et al., 2021). However, targeted therapies to block MMP3 signaling are currently in development. Further, we discovered two targetable genes with current on the market drugs. PSEN1 can be targeted using miR-193a for oligodendrogliomas (Pan et al., 2021). Similarly, KDM1A (LSD1) can be targeted with ladademstat to potentially treat astrocytoma (Lu, et al., 2018; Wang, et al., 2022). PSEN1 and KDM1A (LSD1) have both been associated with cancer previously. These results validate our computational pipeline to identify novel target genes.

**Figure 5.**
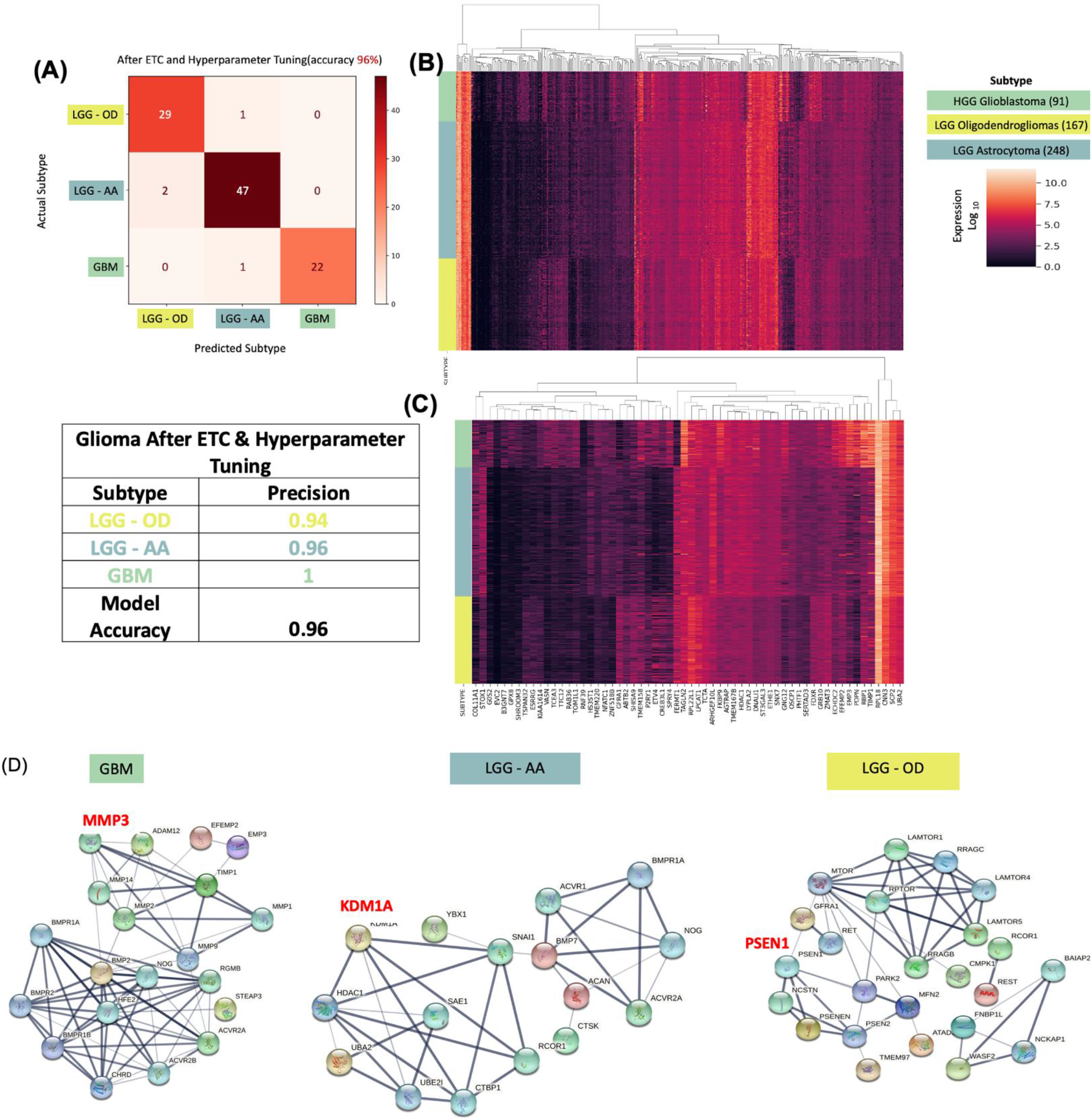
Validation of Computational Pipeline Using Gliomas Dataset identify Distinct Gene Signature and Pathways which are Targetable. A glioma dataset was utilized to validate the computational pipeline. (A) The molecular subtype prediction model had a predictive accuracy of 96%. (B and C) The differential expression analysis was conducted that aided in prioritizing the genes to identify the most impactful pathways. (D) Using StringDB, protein-protein interactions were analyzed. MMP3 can potentially be targeted for glioblastomas (GBM). PSEN1 can potentially be targeted using miR-193a for oligodendrogliomas (LGG-OD) and KDM1A (LSD1) can potentially be targeted using ORY-1001 (ladademstat) to treat astrocytomas (LGG-AA).

## 4. Discussion

Cancer is characterized as a heterogeneous disease genetically and phenotypically with many different subtypes. Rapid diagnosis and accurate identification of cancer subtypes is necessary for disease monitoring and precise treatment options for the patients. For cancer subtyping biomedical researchers have been collecting complex molecular, genetics and clinical data. Integration of these complex omics data demands application of bioinformatics approaches (ref). Recent studies suggest that the application of machine learning (ML) methods could be helpful in expediting these approaches. ML approaches also allows to detect key features from complex datasets reveals their importance. So far a variety of bioinformatics approaches have been applied including Artificial Neural Networks (ANNs), Bayesian Networks (BNs), Support Vector Machines (SVMs) and Decision Trees (DTs) in cancer research for the development of predictive models ((Kourou, et al., 2021). Here we developed a novel machine learning (ML) approach that can predict cancer subtypes high accuracy.

We used TCGA gene expression data to develop a new ML approach to establish gene signature for cancer subtypes. Briefly, we used two different ML approaches to train the model. One model was trained using logistic regression without the use of a feature selection and the other model was trained with the use of a feature selection method, ExtraTreesClassifier (ETC). The overall model accuracy increased by 7 percent (79% to 86%) after implementing ETC for breast cancer subtypes. Further, after conducting hyperparameter tuning the overall predictive accuracy of the model increased to 92%. In order to validate our pipeline, we performed the analysis upon a gliomas dataset (506 patients, 3 subtypes). The model was able to predict the cancer subtype with 96% accuracy. We were able to achieve a subtype predictive accuracy of 96% for gliomas. Thus, our new method that combines logistic regression, hyperparameter tuning, and ExtraTreesClassifier provides high predictive accuracy for cancer subtypes.

Using this approach, we identified that NTRK2, IGF1R, FOXA1, MCM5, and KRT5 are potential targets for breast cancer. These genes have been previously associated with cancers. Additionally, many identified genes have inhibitors currently available commercially, such as linsitinib targeting IGF1R (Murakami, et al., 2016) and entrectinib targeting NTRK2 can possibly be used to inhibit the growth of HER2 Positive (Wang, et al., 2020). FOXA1 can possibly be targeted to inhibit the growth of Triple negative breast cancer (Calissi, et al., 2021). miRNA-214 can be used to target MCM5 and potentially inhibit the growth of Luminal A (Wang, et al., 2022) and miR-601 can be used to target KRT5 to inhibit Luminal B progression (Du et al., 2019). These genes can potentially serve as biomarkers for cancer subtype identification and treatments for personalized therapy. Upon further validation, these targets can lead to new treatment options for aggressive breast cancer subtypes such as Triple negative breast cancer. Similarly, many identified genes, such as KDM1A in astrocytoma, MMP3 (Bufu, et al., 2018) in glioblastoma, and PSEN1 in oligodendroglioma, have previously been correlated with cancer (Feng, et al., 2022; Xu, et al., 2018; Zhang, et al., 2022). These 3 genes also have inhibitors available, such as ladademstat, which can potentially target KDM1A (LSD1) (Salamero, et al., 2020), thus can be used to inhibit the growth of astrocytoma glioma. Similarly, miR-193a can target PSEN1 (Pan, et al., 2021), which can potentially be used to inhibit the growth of oligodendrogliomas. Further, inhibitors for MMP3 are still being tested in cancer (Winer, et al., 2018). In the future, our approach is amenable to incorporate other features such as mutations and drug response and will potentially be applicable to any cancer types. Overall, our approach can also result in an effective and accurate decision making. Even though it is evident that the use of ML methods can improve our understanding of cancer progression, an appropriate level of experimental validation is needed in order for these methods to be considered in the routine clinical practice.

## Data availability

All original data is available through TCGA, and data codes are available through GitHub: https://github.com/vhparikh/TCGA_Pipeline.git.

## Author contribution

EM and VP jointly performed all the analysis, generated pipelines. Both EM and VP wrote the manuscript. EM prepared the first draft. RK provided guidance and resources All authors providing the feedback and edited the manuscript

## Acknowledgement

We thank Dr. Jeremy Goecks for providing the dataset and giving direction and feedback for the project. We also thank the Beaverton Hillsboro Science Expo (BHSE), Northwest Science Expo (NWSE), and International Science and Engineering Fair (ISEF) for giving us valuable feedback and opportunities to present our work.

## References

Alabi, R.O., et al. Deep Machine Learning for Oral Cancer: From Precise Diagnosis to Precision Medicine. Front Oral Health 2021;2:794248.

Arjmand, B., et al. Machine Learning: A New Prospect in Multi-Omics Data Analysis of Cancer. Front Genet 2022;13:824451.

Bac, J., et al. Scikit-Dimension: A Python Package for Intrinsic Dimension Estimation. Entropy (Basel) 2021;23(10).

Bufu, T., et al. Celastrol inhibits colorectal cancer cell proliferation and migration through suppression of MMP3 and MMP7 by the PI3K/AKT signaling pathway. Anticancer Drugs 2018;29(6):530–538.

Cai, Z., et al. Machine learning for multi-omics data integration in cancer. iScience 2022;25(2):103798.

Calissi, G., Lam, E.W. and Link, W. Therapeutic strategies targeting FOXO transcription factors. Nat Rev Drug Discov 2021;20(1):21–38.

Cancer Facts & Figures 2022, A.C.S., https://www.cancer.org/research/cancer-facts-statistics/all-cancer-facts-figures/cancer-facts-figures-2022.html.

Cancer Genome Atlas, N. Comprehensive molecular portraits of human breast tumours. Nature 2012;490(7418):61–70.

Cancer Genome Atlas Research, N., et al. The Cancer Genome Atlas Pan-Cancer analysis project. Nat Genet 2013;45(10):1113–1120.

Chung, M., et al. Early dietitian referral in lung cancer: use of machine learning. BMJ Support Palliat Care 2022.

Ciriello, G., et al. Comprehensive Molecular Portraits of Invasive Lobular Breast Cancer. Cell 2015;163(2):506–519.

Du, H., et al. miR-601 inhibits proliferation, migration and invasion of prostate cancer stem cells by targeting KRT5 to inactivate the Wnt signaling pathway. Int J Clin Exp Pathol 2019;12(12):4361–4379.

Feng, C., Gong, L. and Wang, J. Arborinine from Glycosmis parva leaf extract inhibits clear-cell renal cell carcinoma by inhibiting KDM1A/UBE2O signaling. Food Nutr Res 2022;66.

Fuentes-Baile, M., et al. Differential Effects of IGF-1R Small Molecule Tyrosine Kinase Inhibitors BMS-754807 and OSI-906 on Human Cancer Cell Lines. Cancers (Basel) 2020;12(12).

Guo, Y., et al. Predictors of underutilization of lung cancer screening: a machine learning approach. Eur J Cancer Prev 2022.

Jiang, G., et al. Comprehensive comparison of molecular portraits between cell lines and tumors in breast cancer. BMC Genomics 2016;17 Suppl 7:525.

Kalecky, K., et al. Integrative analysis of breast cancer profiles in TCGA by TNBC subgrouping reveals novel microRNA-specific clusters, including miR-17-92a, distinguishing basal-like 1 and basal-like 2 TNBC subtypes. BMC Cancer 2020;20(1):141.

Kenneth, K.W.T. and Cho, W.C.S. Drug repurposing for cancer therapy in the era of precision medicine. Curr Mol Pharmacol 2022.

Kourou, K., et al. Applied machine learning in cancer research: A systematic review for patient diagnosis, classification and prognosis. Comput Struct Biotechnol J 2021;19:5546–5555.

Krzyszczyk, P., et al. The growing role of precision and personalized medicine for cancer treatment. Technology (Singap World Sci) 2018;6(3-4):79–100.

Lage, I., et al. Efficiently identifying individuals at high risk for treatment resistance in major depressive disorder using electronic health records. J Affect Disord 2022.

Lee, J., et al. Machine learning classifiers-based prediction of normal-tension glaucoma progression in young myopic patients. Jpn J Ophthalmol 2020;64(1):68–76.

Liang, M., et al. Targeting matrix metalloproteinase MMP3 greatly enhances oncolytic virus mediated tumor therapy. Transl Oncol 2021;14(12):101221.

Lu, Z., et al. ORY-1001 Suppresses Cell Growth and Induces Apoptosis in Lung Cancer Through Triggering HK2 Mediated Warburg Effect. Front Pharmacol 2018;9:1411.

Malebary, S.J. and Khan, Y.D. Evaluating machine learning methodologies for identification of cancer driver genes. Sci Rep 2021;11(1):12281.

Manrriquez, E.N., Zakhour, M. and Salani, R. Precision medicine for cervical cancer. Curr Opin Obstet Gynecol 2022;34(1):1–5.

Mateo, J., et al. Delivering precision oncology to patients with cancer. Nat Med 2022;28(4):658–665.

Matplotlib 3.5.1. Matplotlib documentation - Matplotlib 3.5.1 documentation. (n.d.). Retrieved February 17, 2022, from https://matplotlib.org/stable/index.html

Murakami, T., et al. Effective molecular targeting of CDK4/6 and IGF-1R in a rare FUS-ERG fusion CDKN2A-deletion doxorubicin-resistant Ewing’s sarcoma patient-derived orthotopic xenograft (PDOX) nude-mouse model. Oncotarget 2016;7(30):47556–47564.

Niklaus, N.J., et al. The Multifaceted Functions of Autophagy in Breast Cancer Development and Treatment. Cells 2021;10(6).

Pan, X., et al. miR-193a Directly Targets PSEN1 and Inhibits Gastric Cancer Cell Growth, the Activation of PI3K/Akt Signaling Pathway, and the Epithelial-to-Mesenchymal Transition. J Oncol 2021;2021:2804478.

Romero-Cordoba, S.L., et al. Comprehensive omic characterization of breast cancer in Mexican-Hispanic women. Nat Commun 2021;12(1):2245.

Ronquillo, J.G. and Lester, W.T. Precision Medicine Landscape of Genomic Testing for Patients With Cancer in the National Institutes of Health All of Us Database Using Informatics Approaches. JCO Clin Cancer Inform 2022;6:e2100152.

Salamero, O., et al. First-in-Human Phase I Study of Iadademstat (ORY-1001): A First-in-Class Lysine-Specific Histone Demethylase 1A Inhibitor, in Relapsed or Refractory Acute Myeloid Leukemia. J Clin Oncol 2020;38(36):4260–4273.

Szklarczyk, D., et al. STRING v10: protein-protein interaction networks, integrated over the tree of life. Nucleic Acids Res 2015;43(Database issue):D447–452.

Tian, H., et al. Application of Machine Learning Algorithms to Predict Lymph Node Metastasis in Early Gastric Cancer. Front Med (Lausanne) 2021;8:759013.

Tsai, I.J., et al. Machine Learning in Prediction of Bladder Cancer on Clinical Laboratory Data. Diagnostics (Basel) 2022;12(1).

Wang, J., et al. Targeted inhibition of the expression of both MCM5 and MCM7 by miRNA-214 impedes DNA replication and tumorigenesis in hepatocellular carcinoma cells. Cancer Lett 2022;539:215677.

Wang, T., Zhang, F. and Sun, F. ORY-1001, a KDM1A inhibitor, inhibits proliferation, and promotes apoptosis of triple negative breast cancer cells by inactivating androgen receptor. Drug Dev Res 2022;83(1):208–216.

Wang, Y., et al. NTRK Fusions and TRK Inhibitors: Potential Targeted Therapies for Adult Glioblastoma. Front Oncol 2020;10:593578.

Winer, A., Adams, S. and Mignatti, P. Matrix Metalloproteinase Inhibitors in Cancer Therapy: Turning Past Failures Into Future Successes. Mol Cancer Ther 2018;17(6):1147–1155.

Xu, Y., et al. Variants in Notch signalling pathway genes, PSEN1 and MAML2, predict overall survival in Chinese patients with epithelial ovarian cancer. J Cell Mol Med 2018;22(10):4975–4984.

Yuan, H., et al. Application of logistic regression and convolutional neural network in prediction and diagnosis of high-risk populations of lung cancer. Eur J Cancer Prev 2022;31(2):145–151.

Zhang, H., et al. Glioblastoma Treatment Modalities besides Surgery. J Cancer 2019;10(20):4793–4806.

Zhang, K. and Wang, H. [Cancer Genome Atlas Pan-cancer Analysis Project]. Zhongguo Fei Ai Za Zhi 2015;18(4):219–223.

Zhang, W., et al. KDM1A promotes thyroid cancer progression and maintains stemness through the Wnt/beta-catenin signaling pathway. Theranostics 2022;12(4):1500–1517.

